# CRISPR activation of *Tfeb*, a master regulator of autophagy and lysosomal biogenesis, in osteoblast lineage cells increases bone mass and strength

**DOI:** 10.1101/2024.09.26.615175

**Authors:** Alicen James, James Hendrixson, Ilham Kadhim, Adriana Marques-Carvalho, Jacob Laster, Julie Crawford, Jeff Thostenson, Amy Sato, Maria Almeida, Melda Onal

## Abstract

Autophagy is a recycling pathway in which damaged or dysfunctional proteins, protein aggregates, and organelles are delivered to lysosomes for degradation. Insufficiency of autophagy is thought to contribute to several age-related diseases including osteoporosis. Consistent with this, elimination of autophagy from the osteoblast lineage reduces bone formation and causes low bone mass. However, whether increasing autophagy would benefit bone health is unknown. Here, we increased expression of the endogenous Transcription Factor EB gene (*Tfeb*) in osteoblast lineage cells in vivo via CRISPR activation. *Tfeb* overexpression stimulated autophagy and lysosomal biogenesis in osteoblasts. *Tfeb* overexpressing male mice displayed a robust increase in femoral and vertebral cortical thickness at 4.5 months of age. Histomorphometric analysis revealed that the increase in femoral cortical thickness was due to increased bone formation at the periosteal surface. *Tfeb* overexpression also increased femoral trabecular bone volume. Consistent with these results, bone strength was increased in *Tfeb* overexpressing mice. Female *Tfeb* overexpressing mice also displayed a progressive increase in bone mass over time and at 12 months of age had high cortical thickness and trabecular bone volume. This increase in vertebral trabecular bone volume was due to elevated bone formation. Osteoblastic cultures showed that *Tfeb* overexpression increased proliferation and osteoblast formation. Overall, these results demonstrate that stimulation of autophagy in osteoblast lineage cells promotes bone formation and strength and may represent an effective approach to combat osteoporosis.

## INTRODUCTION

Throughout life, bone is remodeled by osteoclasts, which resorb bone, and osteoblasts, which form bone. As osteoblasts form new bone, some become embedded in the bone matrix they produce and differentiate into osteocytes. Osteocytes orchestrate bone remodeling by secreting factors that control bone formation and resorption. When the balance between bone formation and resorption shifts in favor of bone resorption, bone mass is lost. The health and functionality of osteoblast-lineage cells (osteoblast progenitors, osteoblasts, and osteocytes) are essential for bone formation. Therefore, insight into the pathways and mechanisms that control the production, survival, and function of osteoblast lineage cells may identify new therapeutic targets.

Macroautophagy/autophagy is a catabolic process in which cellular contents are engulfed by autophagosomes and delivered to lysosomes for degradation and recycling. Autophagy clears damaged organelles and protein aggregates and turns over cytosolic components. Deletion of genes required for autophagy, such as *Atg7* (autophagy related 7), *Atg5* (autophagy related 5), or *Ulk1* (unc-51 like kinase 1), using various Cre driver strains active in osteoblast lineage cells demonstrates that autophagy is essential for the accrual of bone mass [1-5]. We have shown that deletion of *Atg7* from the entire osteoblast lineage using an Osx1-Cre transgene drastically decreases osteoblast number and bone formation, leading to low bone mass and a 50% fracture rate [2]. Together, these studies demonstrate that autophagy is essential for the maintenance of osteoblast number and bone formation and suggest that insufficient autophagy levels contribute to the decline in bone formation in various skeletal pathologies, such as age-related bone loss [6-8]. However, whether stimulation of autophagy in osteoblast lineage cells could be beneficial for bone in physiological or pathological conditions remains unknown.

To address this question, we sought to stimulate autophagy in osteoblast lineage cells by increasing expression of *Tfeb* (transcription factor EB) – the major transcriptional regulator of genes involved in autophagy and lysosomal biogenesis [9]. This approach is based on evidence that targeted over-expression of *Tfeb* in myotubes [10], neurons [11-13], oligodendrocytes [12], macrophages [14], cardiac myocytes [15], or chondrocytes [16] is sufficient to induce autophagy and alleviate dysfunction in murine models of Pompe disease, Parkinson disease, Alzheimer disease, multiple system atrophy, atherosclerosis, cardiac hypertrophy, and osteoarthritis, respectively. Herein, we used a novel approach, namely in vivo CRISPR activation, to elevate endogenous *Tfeb* expression to stimulate autophagy in osteoblast lineage cells. Stimulation of autophagy greatly increased bone mass by promoting bone formation at both cortical and trabecular bone surfaces.

## RESULTS

### Generation of sgRNA*^Tfeb^* mice

To increase *Tfeb* expression in vivo, we used CRISPR activation (CRISPRa) [17-20]. In this methodology, nuclease-deficient Cas9 (dead Cas9, dCas9), which cannot cut DNA, is fused to one or more transcriptional activator domains (dCas9::activator). To increase transcription of target genes, the dCas9::activator fusion protein is directed to the proximity of the target gene’s transcriptional start site (TSS) via the use of a single guide RNA (sgRNA) that is complementary to that region. The level of transcriptional stimulation varies based on where the dCas9::activator is directed to relative to the TSS of the targeted gene. To identify sgRNAs that facilitate elevated expression of *Tfeb*, we designed 5 sgRNAs targeting the 300 bp region centered on the *Tfeb* TSS with minimal off-target potential (**Figure 1A**). We next tested their activity in vitro using a stromal cell line that expresses dCas9::activator (SP-dCas9-VPR) and identified 3 sgRNAs that elevated *Tfeb* expression 4- to 11-fold in culture (**Figure 1B**). In a previously described murine model, sustained high-level over-expression of *Tfeb* (8-fold compared to controls) in the heart had detrimental effects [21]. To avoid this potential complication, for generation of our mouse model we selected *Tfeb*_sgRNA_4, which stimulates *Tfeb* expression moderately (**Figure 1B**).

**Figure 1.**
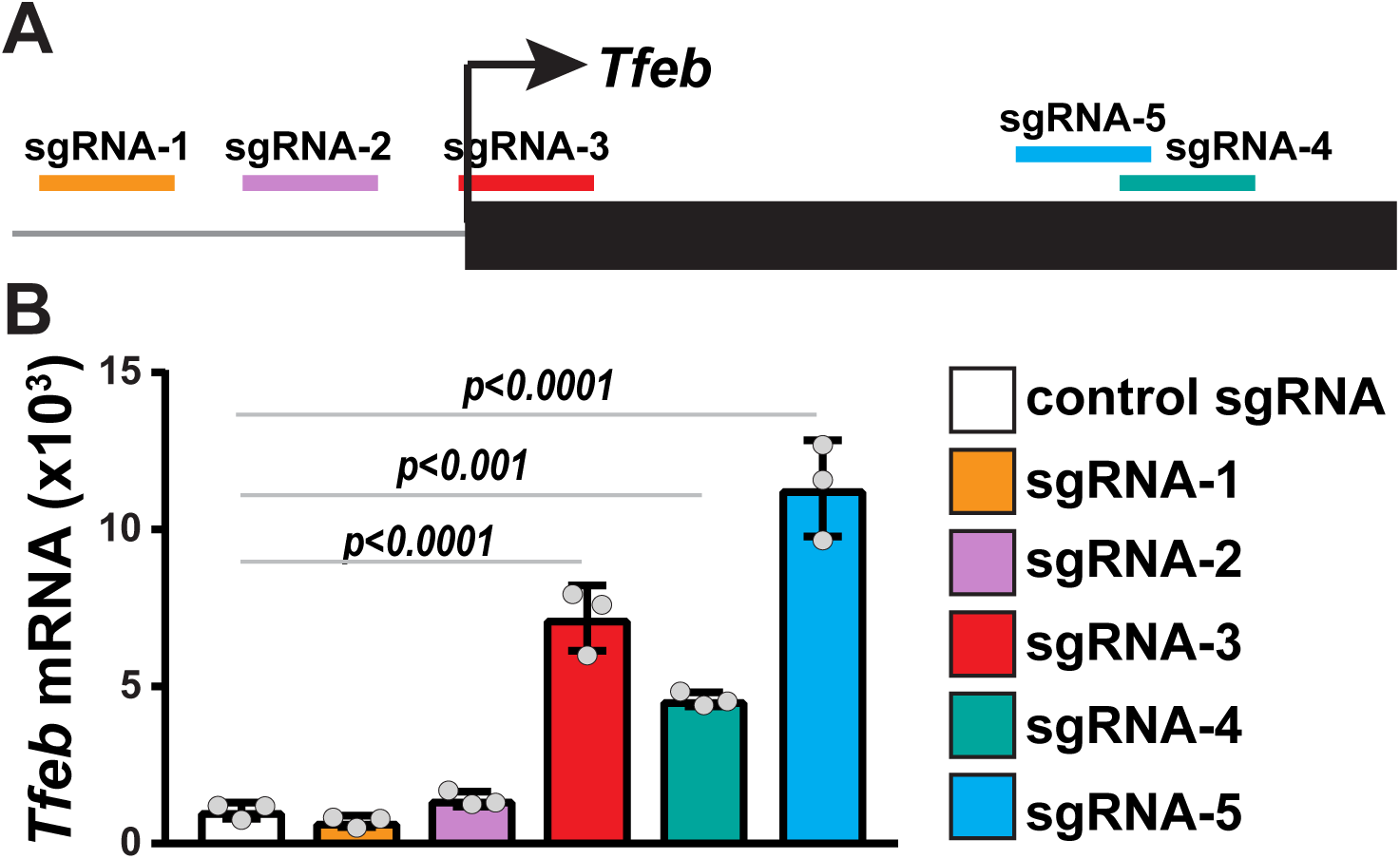
Evaluation of CRISPR activation sgRNAs. (**A**) Diagram showing positions of 5 sgRNAs in relation to the transcription start site (TSS) of *Tfeb* gene. To stimulate transcription of *Tfeb*, the dCas9::SPH/sgRNA complex was targeted to within 150 bp of the TSS on either the template or non-template strand of DNA. (**B**) Gene expression analysis of osteoblastic UAMS-32 cells transfected with dCas9::SPH and sgRNAs targeting *Tfeb*. *Tfeb* mRNA levels were measured by quantitative real-time PCR (qRT-PCR) and normalized to mouse *Actb*. Data are presented as mean ± standard deviation (SD), n=3 wells per group. *p* values are calculated by one-way ANOVA followed by Dunnett’s multiple comparison test comparing each to the control group.

To obtain ubiquitous and consistent expression of the selected sgRNA for CRISPRa, we inserted an expression cassette encoding it into a safe harbor locus. In previous studies, we successfully used this approach to express a sgRNA for in vivo CRISPR interference [22]. Similarly, herein we introduced a cassette expressing *Tfeb*_sgRNA_4 into the murine *Rosa26* locus, producing sgRNA*^Tfeb^* mice. As expected, hemizygous or homozygous sgRNA*^Tfeb^* mice were born at expected Mendelian ratios and were grossly indistinguishable from wild-type mice.

### *Tfeb* elevation in the osteoblast lineage stimulates autophagy and lysosomal biogenesis

To increase endogenous *Tfeb* in osteoblast-lineage cells in vivo, we crossed sgRNA*^Tfeb^* mice with CRa [19] and Osx1-Cre mice [23] (**Figure 2A**). CRa transgenic mice contain a floxed stop cassette that prevents expression of dCas9::activator (dCas9::SunTag-p65-HSF1 or dCas9::SPH) until activated by a Cre driver strain [19]. In triple transgenic CRa;Osx1-Cre;sgRNA*^Tfeb^* mice, from here on referred to as *Tfeb*^CRa^ mice, dCas9::SPH is expressed only in cells targeted by Osx1-Cre. *Tfeb*^CRa^ mice showed a ∼1.4-fold increase in *Tfeb* mRNA in vertebral bones compared to controls (**Figure 2B**). A 2.7-fold increase in *Tfeb* mRNA was also seen in calvaria and this was associated with increased expression of several genes involved in lysosomal biogenesis and autophagy (**Figure 2C**).

**Figure 2.**
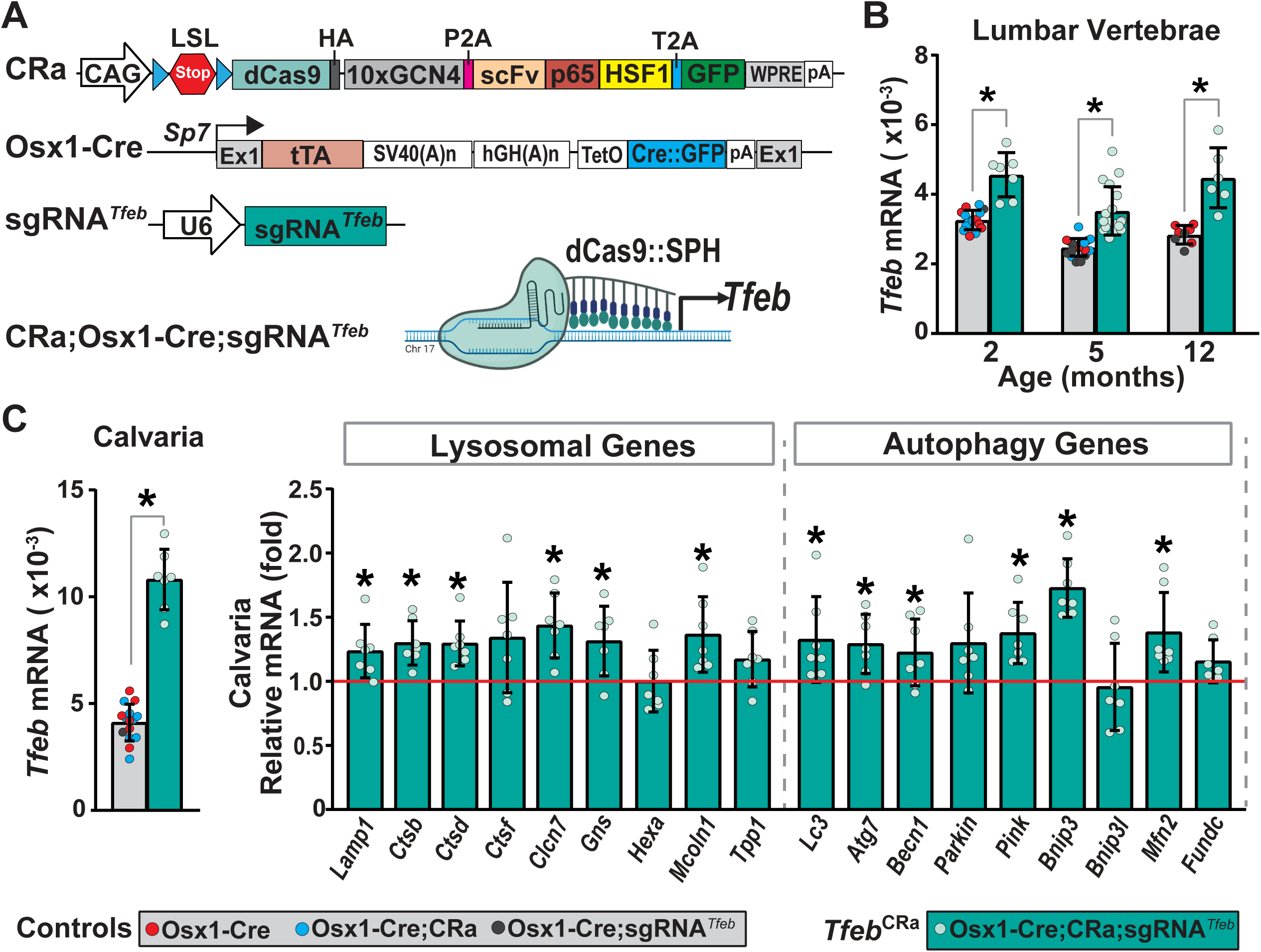
Elevation of *Tfeb* in osteoblast lineage cells via CRISPR activation stimulates expression of genes involved in autophagy- and lysosomal biogenesis. (**A**) Transgene design of murine models used for CRISPR activation. Specifically, CRISPR-dCas9-activator (CRa), Osx1-Cre and sgRNA*^Tfeb^* mice were crossed to obtain triple transgenic CRa;Osx1-Cre;sgRNA*^Tfeb^* (*Tfeb*^CRa^) mice. (**B**) *Tfeb* mRNA levels were measured in lumbar vertebrae 5 (L5) of *Tfeb*^CRa^ mice and their littermate Cre controls by quantitative real-time PCR (qRT-PCR) at 2, 5, or 12 months of age. (**C**) mRNA levels of *Tfeb* (left) and genes involved in lysosomal biogenesis and autophagy (right) were measured in calvaria of 2-month-old *Tfeb*^CRa^ mice and their littermate controls via qRT-PCR. For all qRT-PCR analyses, mRNA levels were normalized to mouse *Actb*. Lysosome- and autophagy-related gene expression is represented as fold change over Cre-controls. Bars indicate mean ± SD. n = 7-15 mice/group. *, *p* < 0.05 as calculated by unpaired *t*-test. Of note, unpaired *t*-test with Welch’s correction is used in cases where the SD of groups were different.

To examine whether *Tfeb* over-expression stimulates autophagy in osteoblast lineage cells, we cultured stromal cells isolated from femurs of *Tfeb*^CRa^ mice and littermate controls in osteogenic differentiation medium for 3 weeks. Western blot analysis confirmed that *Tfeb*^CRa^ osteoblasts had higher TFEB levels compared to Cre controls (**Figure 3A**). To quantify autophagic flux, the osteoblasts were treated with Bafilomycin, which blocks lysosomal degradation, or vehicle. Degradation of LC3-II (microtubule-associated protein 1 light chain 3 alpha - II) and SQSTM1/p62 (sequestosome 1) are commonly used to measure autophagic flux. As the autophagosome forms, LC3 is lipidated and incorporated into the autophagosome as LC3-II. LC3-II interacts with receptor proteins which target various cytoplasmic components for autophagic degradation. p62 is one such autophagic receptor protein that is engulfed by the autophagosome along with its cargo. As such, LC3-II and p62 are degraded by the lysosome when the autophagosome fuses with the lysosome during the autophagy process. Therefore, the difference in the level of these proteins between vehicle- and Bafilomycin-treated cells provides a measure of autophagic flux. In *Tfeb*^CRa^ osteoblasts, elevated TFEB was associated with increased autophagic flux as indicated by increased accumulation of LC3-II and p62 following treatment with Bafilomycin (**Figure 3A**). An increase in lysosomal biogenesis was indicated by LAMP1 (lysosomal-associated membrane protein 1) protein levels and LysoTracker staining (**Figure 3A-B**). Taken together, these results demonstrate that the elevation of endogenous *Tfeb* in osteoblast lineage cells of *Tfeb*^CRa^ mice is sufficient to stimulate autophagy and lysosomal biogenesis.

**Figure 3.**
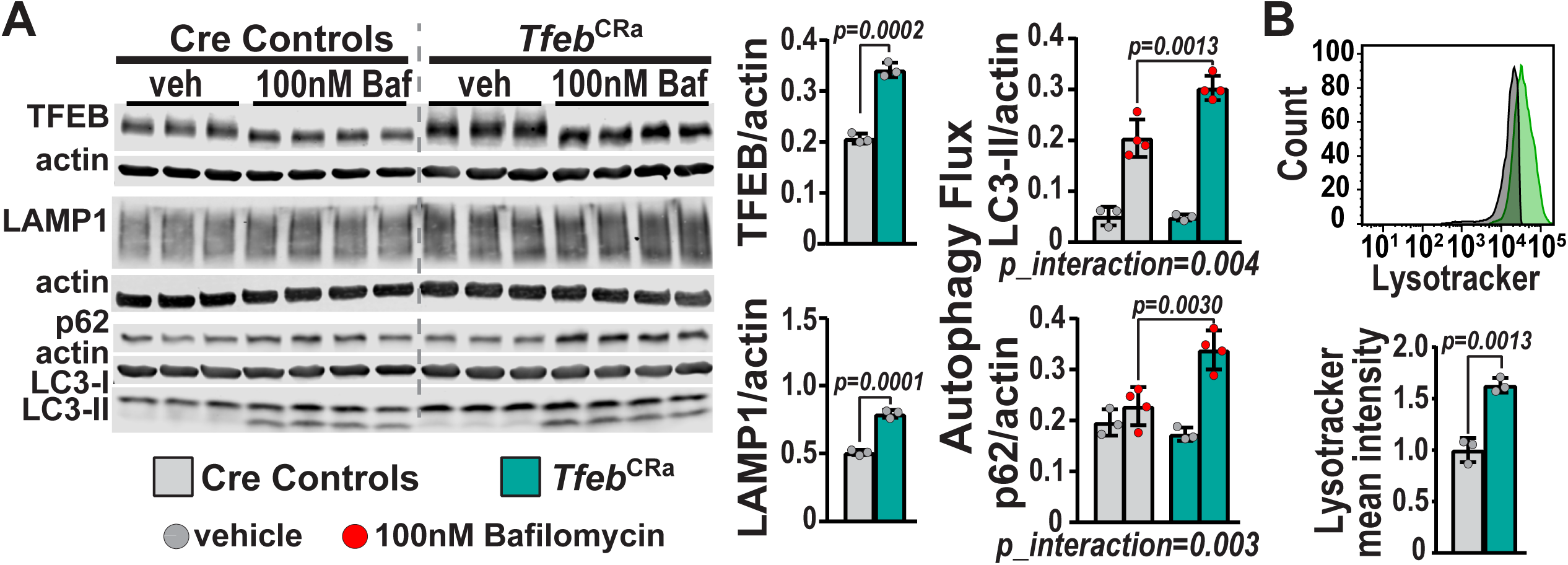
Elevation of TFEB in osteoblasts induces autophagy and lysosomal biogenesis. (**A**) Bone marrow stromal cells were isolated from femurs and tibias of 5-month-old female *Tfeb*^CRa^ mice and littermate Cre controls, and cultured in osteogenic media for 21 days. Cells were treated with vehicle (PBS) or 100 nM Bafilomycin for 6 hours. Western blot analysis was performed to measure TFEB, LAMP1, p62, actin, and LC3 levels. For quantification, protein levels were normalized to actin levels. n = 3-4 wells per group. (**B)** Bone marrow stromal cells were isolated from femurs and tibias of 5-month-old female *Tfeb*^CRa^ mice and littermate Cre controls, and cultured in osteogenic media until cells reached 80% confluency. Lysotracker mean intensity was measured by flow cytometry. n = 3 wells/group. Bars indicate mean ± SD. Indicated *p* values were calculated by unpaired *t*-test or 2-way ANOVA followed by Tukey’s post-hoc analysis.

### Elevation of *Tfeb* in the osteoblast lineage increases bone mass and strength

Mice carrying the Osx1-Cre transgene are known to be smaller and have low bone mass when compared to mice without this transgene [24]. To minimize the skeletal phenotypes associated with the Osx1-Cre transgene, we utilized its doxycycline regulation [23,24]. In the Osx1-Cre transgene, Cre expression is under the control of a tetracycline-controlled transcriptional activator (tTA in the Tet-off system) which allows temporal regulation of Cre expression via administration of tetracycline or doxycycline. Therefore, we maintained breeders and their progeny (up to 3 weeks of age) on a doxycycline-containing diet (**Figure 4A**, **Figure 4-7**), which prevents Cre expression and bypasses the impact of the Osx1-Cre transgene on skeletal development [25]. Nonetheless, to account for any effects of the Osx1-Cre transgene as mice age, we compared *Tfeb*^CRa^ mice to their Cre-harboring littermates.

**Figure 4.**
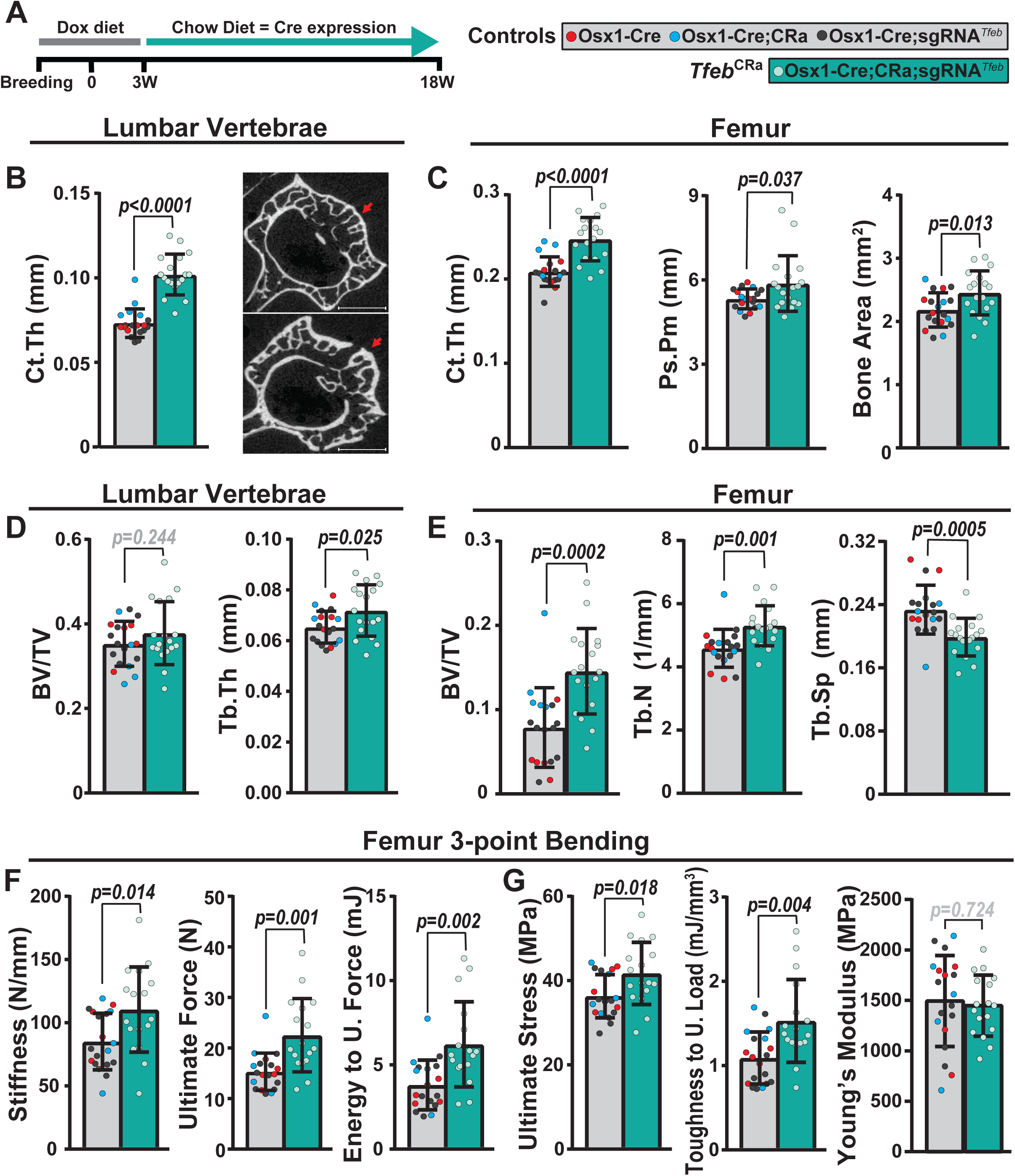
*Tfeb* elevation in the osteoblast lineage increases bone mass and strength. (**A**) Mice remained on a doxycycline-containing diet until 3 weeks of age to repress expression of the Osx1-Cre transgene. At 3 weeks, mice were switched to a chow diet, allowing expression of Osx1-Cre. (**B-E**) µCT analysis was performed on lumbar vertebrae 4 (L4) and femurs from 4.5-month-old male *Tfeb*^CRa^ mice and their littermate controls. (**B**) Cortical thickness (Ct.Th) was measured in L4 (left), representative image (right). (**C**) Ct.Th, periosteal perimeter (Ps.Pm), and bone area were measured at the femoral midshaft. (**D**) Trabecular bone volume over tissue volume (BV/TV) and trabecular thickness (Tb.Th) were measured in L4. (**E**) BV/TV, Tb.Th, and trabecular spacing (Tb.Sp) were measured at the distal femur. (**F** and **G**) Femoral 3-point-bending biomechanical testing was performed on 4.5-month-old male *Tfeb*^CRa^ mice and their littermate controls to assess extrinsic (**F**) and intrinsic (**G**) properties of bone. Bars indicate mean ± SD. n = 17-19 mice/group. Indicated *p* values were calculated by unpaired *t*-test with Welch’s correction.

We first examined a cohort of 4.5- to 5-month-old mice, an age at which growth is completed and peak bone mass is attained. MicroCT analysis of male *Tfeb*^CRa^ mice revealed higher cortical thickness in both the spine (∼39% increase) and femur (∼18% increase) (**Figure 4B-C**). The change in cortical thickness in femur was due to increased periosteal perimeter (**Figure 4C**), while no changes were detected in the endosteal perimeter (**Figure S1A**). *Tfeb* over-expression in the osteoblast lineage also increased vertebral trabecular thickness without a significant impact on trabecular bone volume (**Figure 4D**). In contrast, trabecular bone in the distal femur was increased due to higher trabecular number (**Figure 4E**). We observed a similar phenotype in female *Tfeb*^CRa^ mice (**Figure S2**). These results demonstrate that *Tfeb* over-expression in osteoblast lineage cells is beneficial for both trabecular and cortical compartments in young adult mice.

We next tested whether the increases in bone mass of *Tfeb*^CRa^ mice improved biomechanical properties by performing 3-point bending testing of femurs. This analysis revealed that both extrinsic/structural (stiffness, ultimate force, and energy to ultimate force) and the intrinsic/material properties of the bone (ultimate stress and toughness to ultimate load) were increased in *Tfeb*^CRa^ mice compared to controls (**Figure 4F-G**). However, there were no differences in the Young’s Modulus (**Figure 4G**), which is calculated as stiffness normalized by the tissue distribution and geometry, indicating that the gains in stiffness are due to beneficial changes in bone geometry and microarchitecture, as opposed to alterations in tissue properties (degree of mineralization or collagen crosslinking). These results demonstrate that *Tfeb* over-expression increases the amount of bone, which is of normal quality, resulting in stronger bones.

### Elevation of *Tfeb* in the osteoblast lineage increases bone formation

To identify the cellular mechanisms underpinning the high bone mass phenotype of *Tfeb*^CRa^ mice, we performed dynamic histomorphometry of femurs from male *Tfeb*^CRa^ and Cre control mice. At the periosteal surface, the mineralizing surface, mineral apposition rate, and bone formation rate of *Tfeb*^CRa^ mice were higher than Cre controls (**Figure 5A-B**). The femoral endosteal bone formation was similar in *Tfeb*^CRa^ mice and controls (**Figure S3A**). These results indicate that the increase in femoral cortical thickness of *Tfeb*^CRa^ mice was due to higher periosteal bone formation. On the other hand, the increase in femoral trabecular bone volume of *Tfeb*^CRa^ mice was associated with increased mineral apposition rate at the trabecular surface (**Figure 5C** and **Figure S3B**). Next, we measured expression of osteoblast and osteoclast marker genes in mRNA isolated from lumbar vertebrae. This analysis revealed elevated levels of both osteoblast (*Sp7* - *Sp7 transcription factor 7/Osterix* and *Col1a1-collagen, type I, alpha 1*) and osteoclast (*Acp5 - acid phosphatase 5, tartrate resistant* and *CtsK - Cathepsin K*) marker genes (**Figure 5D-E**), suggesting a high turnover state that favors bone formation which results in a modest increase vertebral trabecular thickness (**Figure 4D**).

**Figure 5.**
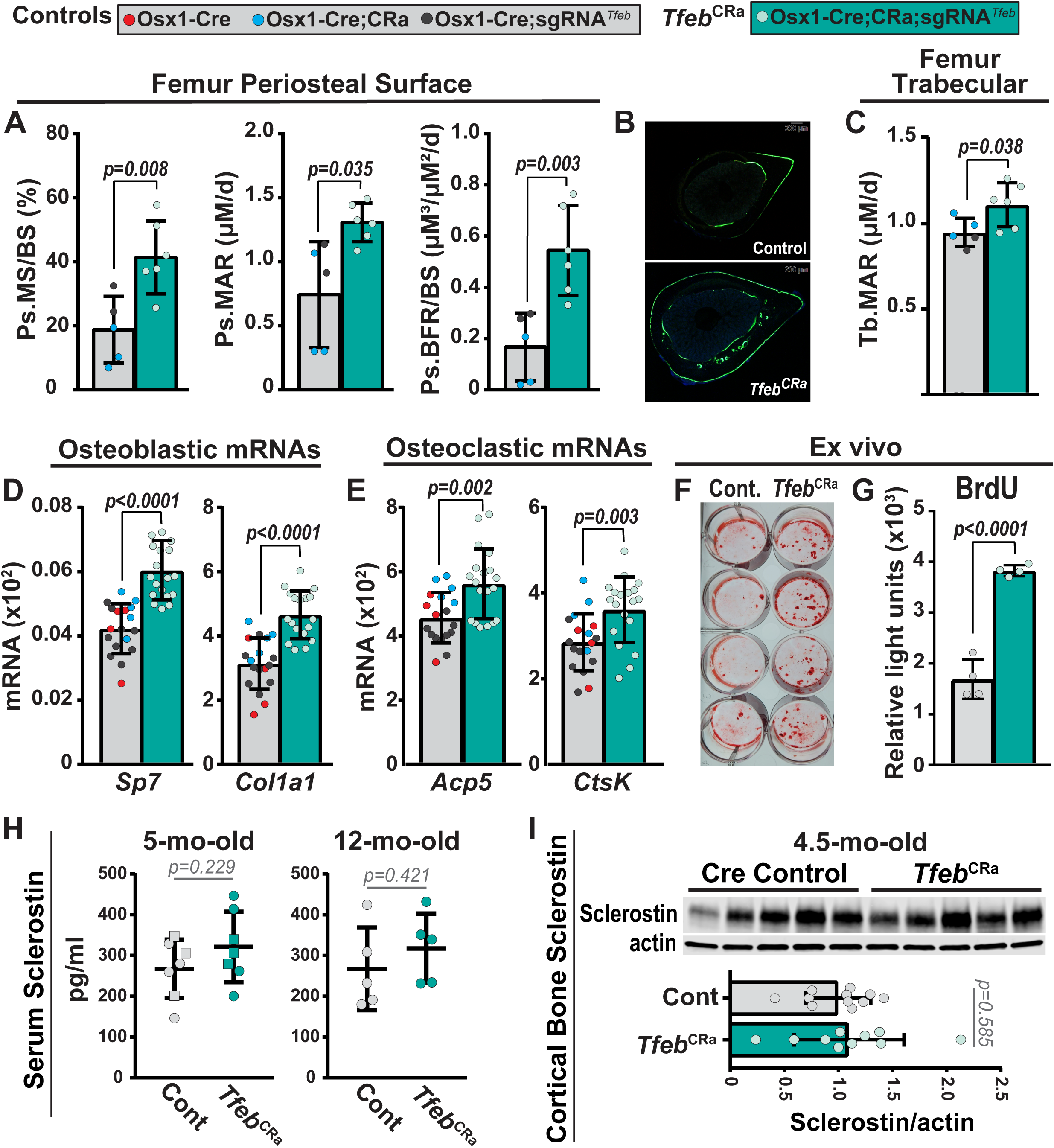
*Tfeb* elevation in the osteoblast lineage increases bone formation. (**A-C**) Dynamic histomorphometry was performed on femurs of 4.5-month-old male *Tfeb*^CRa^ mice and their littermate Cre controls. n = 5-6 mice/group. (**A**) Quantification of the mineralizing surface per bone surface (MS/BS), mineral apposition rate (MAR), and bone formation rate per bone surface (BFR/BS) at the periosteal surface. (**B**) Representative image of a histological cross section at the femoral diaphysis taken using a GFP filter. (**C**) Trabecular MAR (Tb. MAR). (**D** and **E**) Gene expression analysis of 4.5-month-old male *Tfeb*^CRa^ mice and their littermate controls. n = 18-19 mice/group. *Sp7* and *Col1a1* (**D**) and *Acp5* and *CtsK* (**E**) mRNA levels were measured in lumbar vertebrae 5 (L5) of *Tfeb*^CRa^ mice and their littermate controls by qRT-PCR. For all qRT-PCR analyses, mRNA levels were normalized to mouse *Actb*. (**F** and **G**) Bone marrow stromal cells were isolated from femurs and tibias of 5-month-old female *Tfeb*^CRa^ mice and littermate Cre controls, and cultured in osteogenic media. n = 4 wells/group. (**F**) Alizarin red stain of mineral apposition. (**G**) BrdU analysis of proliferating cells. (**H**) Circulating sclerostin levels of 5- and 12-month (mo)-old male (square) and female (circle) *Tfeb*^CRa^ and control mice were measured using ELISA. n=5-7 mice/group. (**I**) Sclerostin and actin levels were measured in cortical bone preparations (humeri shafts) of *Tfeb*^CRa^ and control mice via western blot analysis. The bar graph indicates sclerostin levels normalized to actin levels. n= 10-11 mice/group. Bars indicate mean ± SD. Indicated *p* values were calculated by unpaired *t*-test. Unpaired *t*-test with Welch’s correction was used if the SD of groups were different.

In line with these in vivo findings, *Tfeb* over-expression increased mineral deposition and proliferation in cultured osteoblastic cells (**Figure 5F-G**). Sclerostin is a Wnt signaling inhibitor that is produced mainly by osteocytes to inhibit bone formation. Previous work has suggested that sclerostin is degraded in the lysosomes [26]. This raises the possibility that sclerostin levels could be altered in *Tfeb*^CRa^ mice. We examined this possible mechanism by quantifying sclerostin levels in serum and cortical bone, but did not detect any differences between *Tfeb*^CRa^ and Cre-control mice (**Figure 5H-I**). Together these results suggest that elevation of *Tfeb* in Osx1-Cre targeted cells increases bone formation likely by promoting the proliferation of osteoblast lineage cells.

### Elevation of *Tfeb* in osteoblast lineage cells increases bone mass up to 12 months of age

To gain a better idea of the progression of the bone phenotype, we next performed serial analysis of bone mineral density (BMD) in live mice using dual-energy x-ray absorptiometry (DXA) from 3 to 12 months of age. *Tfeb*^CRa^ mice exhibited high spine and femur BMD beginning at 3 and 7 months, respectively, that remained high up to the time of sacrifice at 12 months (**Figure 6A-B**). To address if elevation of *Tfeb* increased bone mass post-growth, we compared BMDs at 5, 9, and 12 months of age and showed that the high bone mass phenotype became more pronounced up to 9 months of age and remained elevated up to 12 months of age (**Figure 6B-C**). Body weights of *Tfeb*^CRa^ and Cre control mice were similar at all ages.

**Figure 6.**
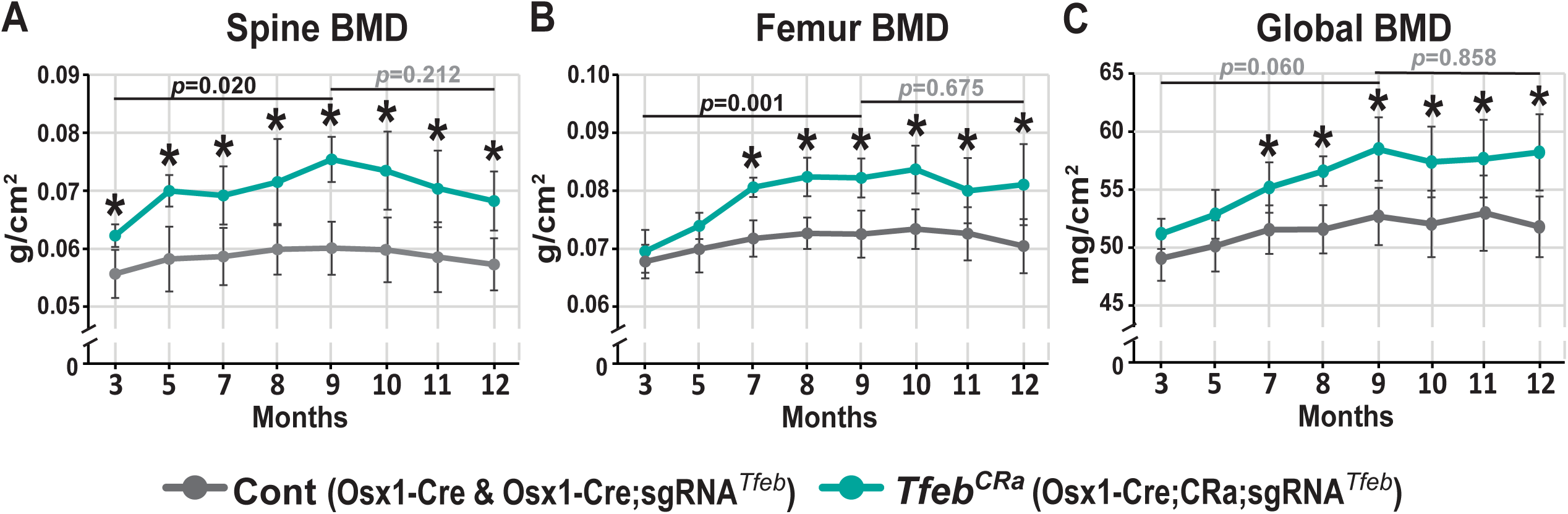
*Tfeb* elevation in the osteoblast lineage progressively increases bone mass up to 12 months of age. (**A-C**) Serial vertebral (**A**), femoral (**B**) and whole body (**C**) BMD measurements were collected via DXA analysis in female *Tfeb*^CRa^ mice and their littermate Cre controls from 3-12 months of age. Bars indicate mean ± SD. n = 5-8 mice/group. *, *p* < 0.05 as calculated by unpaired *t*-test comparing the *Tfeb*^CRa^ mice and their littermate Cre controls at each time point. Specifically indicated *p* values were calculated by repeated measures model (detailed in the methods section) comparing the difference in genotypes at 5 vs. 9 months; or 5 vs. 12 months.

MicroCT analysis performed at 12 months of age showed that *Tfeb*^CRa^ mice maintained elevated cortical thickness in both the spine and femur (**Figure 7A**). At 12 months of age, *Tfeb*^CRa^ mice had elevated vertebral bone volume compared to Cre-controls (**Figure 7B** and **Figure S4A**). Of note, at 5 months of age vertebral trabecular bone volume of female *Tfeb*^CRa^ mice was similar to Cre controls (**Figure S2C**), suggesting that elevated *Tfeb* causes a progressive increase in vertebral trabecular bone volume. Mice lose femoral trabecular bone before age-related bone loss occurs in other sites or compartments [27-29]. Specifically, by the time mice are 12 months of age, they have already lost most of their femoral trabecular bone [27-29]. Consistent with this, Cre control mice had less than 4% trabecular bone volume at 12 months of age (**Figure 7B**). In contrast, at this age *Tfeb*^CRa^ mice had approximately 40% trabecular bone volume, demonstrating that *Tfeb*^CRa^ mice retained their femoral trabecular bone volume up to 12 months of age (**Figure 7B**).

**Figure 7.**
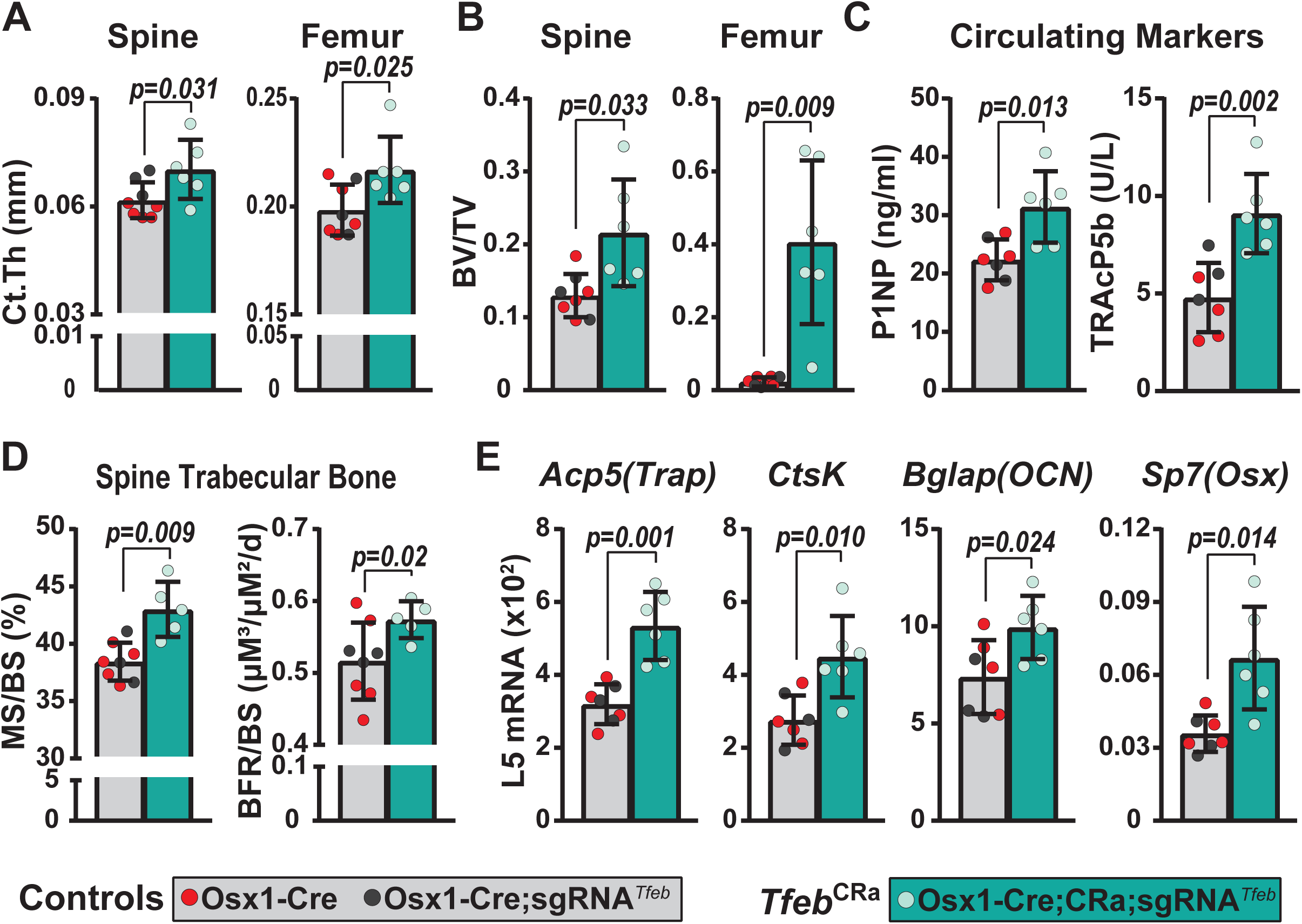
*Tfeb* elevation in the osteoblast lineage remains anabolic at 12 months of age. (**A** and **B**) µCT analysis was performed on lumbar vertebrae 4 (L4) and femurs from 12-month-old female *Tfeb*^CRa^ mice and their littermate controls. (**A**) Cortical thickness (Ct.Th) was measured in L4 (spine) and at the femoral midshaft (femur). (**B**) Trabecular bone volume over tissue volume (BV/TV) was measured in L4 (spine) and at the femoral distal metaphysis (femur). (**C**) Quantification of P1NP, a circulating bone formation marker, and TRAcP5b, a circulating marker of bone resorption, via ELISA. (**D**) Quantification of the mineralizing surface per bone surface (MS/BS) and bone formation rate per bone surface (BFR/BS) at the vertebral trabecular bone. (**E**) *Acp5*, *CtsK*, *Bglap*, and *Sp7* mRNA levels were measured in lumbar vertebrae 5 (L5) of *Tfeb*^CRa^ mice and their littermate controls by qRT-PCR. mRNA levels were normalized to mouse *Actb*. Bars indicate mean ± SD. n = 5-8 mice/group. Indicated *p* values were calculated by unpaired *t*-test. Unpaired *t*-test with Welch’s correction was used if the SD of groups were different.

### Elevation of *Tfeb* in the osteoblast lineage is anabolic up to 12 months of age

We next sought to examine the cellular basis of the high bone mass phenotype of *Tfeb*^CRa^ mice at 12 months of age. Circulating bone formation and resorption markers were both elevated in the serum of *Tfeb*^CRa^ mice, suggesting a high turnover state (**Figure 7C**). However, as high bone volume may contribute to elevation of circulating markers, we performed further analysis of bone formation and resorption.

Dynamic histomorphometry in femurs revealed that at 12 months of age, periosteal and endosteal bone formation of *Tfeb*^CRa^ and Cre-control mice were similar (**Figure S4C-D**). This suggests that while *Tfeb*^CRa^ mice maintain elevated cortical thickness and periosteal perimeter compared to controls, periosteal bone apposition does not remain elevated. As control mice do not have femoral trabecular bone at this age, a comparison of trabecular bone formation in femurs is not possible. However, as can be seen in representative images shown in **Figure S5**, bone formation is ongoing at the femoral trabecular compartment in *Tfeb*^CRa^ mice.

At the vertebral trabecular bone compartment, the mineralizing surface and bone formation rate of 12-month-old *Tfeb*^CRa^ mice were higher compared to Cre controls (**Figure 7D**). Consistent with this, expression of osteoblast marker genes were elevated in lumbar vertebrae of *Tfeb*^CRa^ mice (**Figure 7E**). While we did not measure osteoclast numbers at this age, expression of osteoclast marker genes was also elevated in lumbar vertebrae (**Figure 7E**), suggesting that the higher bone volume of *Tfeb*^CRa^ mice is not due to decreased bone resorption. Together, these results suggest that *Tfeb* over-expression in the osteoblast lineage increases bone formation to cause a progressive increase in the vertebral trabecular bone volume of *Tfeb*^CRa^ mice.

### Elevation of *Tfeb* in osteoblast lineage cells increases bone mass during growth

Having established that *Tfeb* over-expression in osteoblast lineage cells is anabolic post-growth (beyond 5 months of age) (**Figure 6**), we next examined whether *Tfeb* over-expression could alter bone mass accrual during development and growth. To address this question, we allowed continuous expression of Osx1-Cre transgene in breeders and pups by maintaining all mice on a chow diet. We then compared a combination of wild-type, CRa, and sgRNA mice with or without the Osx1-Cre transgene (used as no-Cre and Cre controls) with the *Tfeb*^CRa^ mice at 8 weeks of age. As previously reported [24], mice carrying the Osx1-Cre transgene were smaller and had low bone mass when compared to mice without Osx1-Cre transgene (**Figure 8A**). Therefore, for detailed microCT analysis, we only used mice bearing the Osx1-Cre transgene. At this age, *Tfeb* over-expression increased vertebral, but not femoral, cortical thickness (**Figure 8B**). Moreover, elevated *Tfeb* increased femoral trabecular bone volume, but did not alter vertebral trabecular bone volume (**Figure 8C** and **Figure S5**). These results suggest that *Tfeb* over-expression in the osteoblast lineage promotes bone mass accrual during growth.

**Figure 8.**
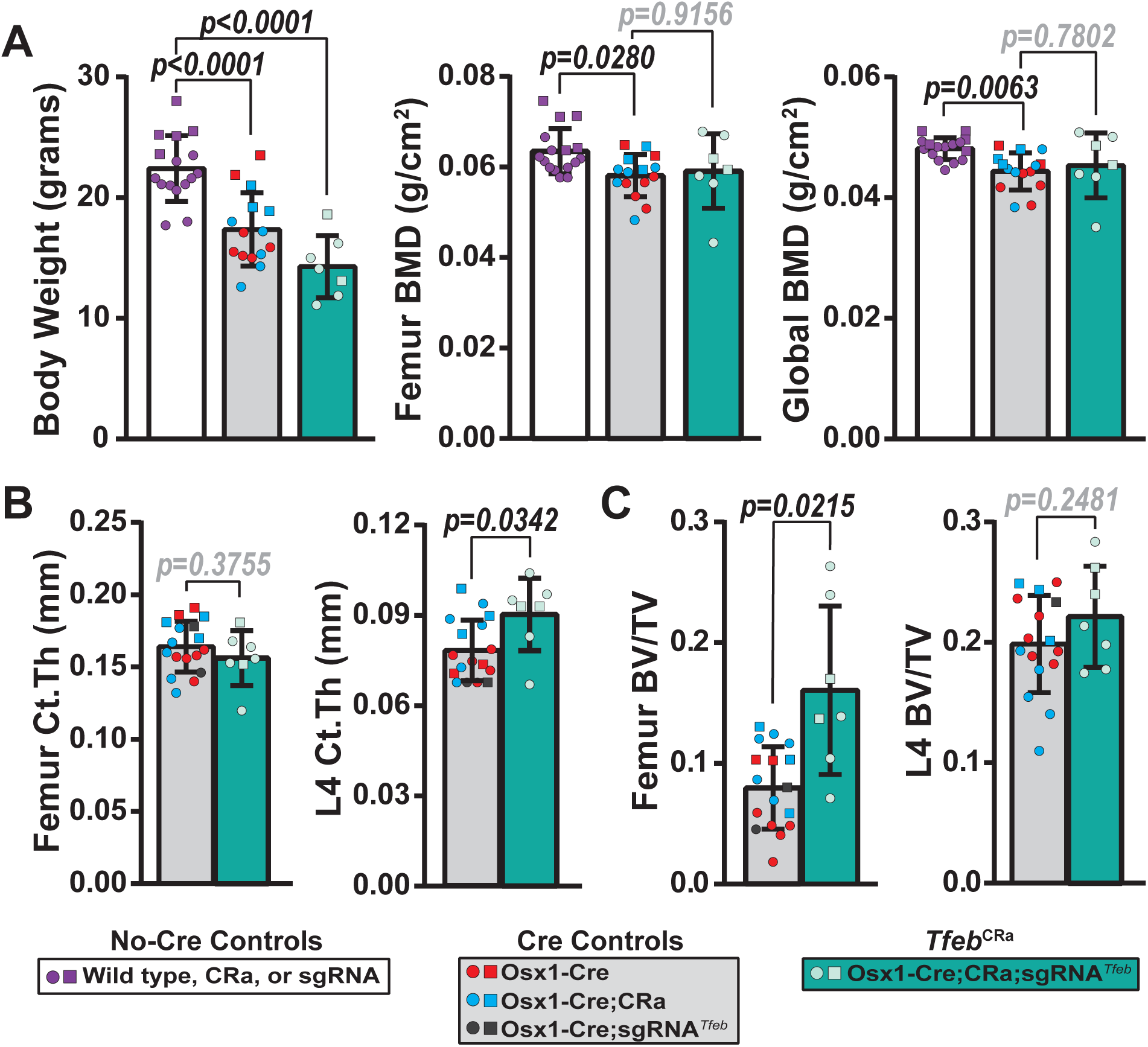
Elevation of *Tfeb* in osteoblast-lineage cells increases bone mass during growth. (**A**) Bone mineral density (BMD) was measured via dual-energy x-ray absorptiometry (DXA) in male (square) and female (circle) *Tfeb*^CRa^ mice and their littermate No-Cre and Cre controls at 2 months of age. (**B** and **C**) µCT analysis was performed on femurs and lumbar vertebrae 4 (L4) of 2-month-old male (square) and female (circle) *Tfeb*^CRa^ mice and their littermate Cre controls. (**B**) Cortical thickness (Ct.Th) was measured at the femoral midshaft and L4. (**C**) Bone volume over tissue volume (BV/TV) was measured at distal femur and L4. Bars indicate mean ± SD. n = 7-17 mice/group. Indicated *p* values were calculated by One-way ANOVA followed by Tukey’s post-hoc analysis (**A**) or unpaired *t*-test (**B,C**).

## DISCUSSION

We have previously shown that the elimination of autophagy from osteoblast-lineage cells using an Osx1-Cre transgene decreases bone turnover, decreases trabecular bone volume, and reduces cortical thickness by decreasing periosteal bone formation [2]. This work, along with others [1-5], establishes that autophagy in the osteoblast lineage is essential for bone formation. However, whether stimulation of autophagy would have a beneficial impact on bone was unknown. In the present study, we aimed to address this question by stimulating autophagy via elevating the expression of endogenous *Tfeb*, a master transcriptional regulator of autophagy and lysosomal biogenesis, in the entire osteoblast lineage. We confirmed that elevation of *Tfeb* in osteoblast lineage cells increases the autophagic flux of osteoblasts. Moreover, we showed that elevation of *Tfeb* in the osteoblast lineage results in a skeletal and cellular phenotype that is the opposite of autophagy elimination in this lineage [2]. Specifically, *Tfeb*^CRa^ mice have increased bone turnover, increased trabecular bone volume, and increased cortical thickness due to increased periosteal bone formation. Based on these results, we conclude that the anabolic impact of *Tfeb* elevation in the osteoblast lineage is due to stimulation of autophagy.

We used CRISPR activation (CRISPRa) for our gain-of-function studies. This new methodology has various benefits over traditional transgene-based over-expression approaches. In traditional approaches, transgene expression is influenced by copy number, integrity, and insertion site into the genome [30-33]. Transgenic approaches can also lead to undesirably high expression levels causing unwanted changes in transcription [21]. In contrast, CRISPRa stimulates the transcription of endogenous genes, which avoids the copy number and integration site effects. CRISPRa also provides control over the magnitude of gene stimulation. Herein, we show that expression of a sgRNA from a safe harbor locus is sufficient to elevate target gene expression in vivo via CRISPR activation. In traditional transgenic approaches, transgene expression may also be suppressed over time [30-32]. In contrast, herein we show that transcriptional stimulation via CRISPRa persists up to at least 12 months of age. Overall, our results demonstrate that CRISPRa can effectively and reliably stimulate target genes in a cell type-specific manner in mice.

Our ex vivo studies suggest that the anabolic impact of *Tfeb* over-expression is due to elevated proliferation of osteoblast progenitors. In recent years, different groups have identified novel osteoblast progenitor populations using combinations of scRNA-seq and lineage tracing [34-36]. However, there is not yet a consensus about the exact identity or function of these different cell populations. Therefore, future studies are needed to examine the contribution of precursor cell populations to the anabolism observed in *Tfeb*^CRa^ mice. In addition, recent studies using single-cell RNA sequencing (scRNA-seq) show that the Osx1-Cre transgene used in our studies targets several different cell populations at the periosteum as well as at the endosteal and trabecular surfaces, including but not limited to osteoblast precursors, osteoblasts, and osteocytes [37]. Therefore, future scRNA-seq studies complemented by functional studies will be necessary to identify the cell types and molecular processes affected by *Tfeb* over-expression in *Tfeb*^CRa^ mice.

In non-osseous cell types, genetic stimulation of autophagy in young mice decreases oxidative stress [38], improves muscle strength and motor performance [39], dampens cytokine production [14,39,40], clears protein aggregates [12,14], decreases ER stress [40,41] and prevents apoptosis [14]. The majority of these effects were observed in murine disease models that use young mice or in response to pathological challenges such as a high-fat diet. Herein, we show that elevation of autophagy in osteoblast lineage cells promotes bone accrual during growth and adulthood under physiological conditions. While further studies are required to examine the molecular mechanisms underpinning this effect, our murine model will be useful for determining the therapeutic potential of autophagy stimulation in various skeletal pathologies. For instance, previous studies suggest that macroautophagy maintains protein homeostasis in osteoblasts by clearing misfolded collagen (10, 29). In osteogenesis imperfecta (OI), mutations exaggerate collagen misfolding beyond the clearance capacity of cells, resulting in aggregation of misfolded collagen in the ER [42-44]. This loss of proteostasis in OI is thought to cause osteoblast dysfunction and contribute to bone pathology. Accordingly, a previous study used rapamycin – an mTOR inhibitor – to assess if elevation of autophagy improved OI pathology [45]. Treatment of growing OI mice with this mTOR inhibitor had opposing effects on trabecular versus cortical compartments. Overall, this treatment did not improve the strength of OI bones. However, as the authors point out, this study was not able to distinguish between the effects of autophagy and mTOR signaling (which mediates cell growth, energy metabolism, and protein synthesis), or which cell types were affected by rapamycin. Our *Tfeb*^CRa^ model provides improved molecular and cellular specificity, and, therefore, represents a valuable tool for determining if stimulating autophagy can relieve ER stress in OI osteoblasts and ameliorate OI pathology.

Dysfunctional autophagy is a hallmark of aging [46]. Several studies have examined the impact of autophagy stimulation in the setting of aging or age-related pathologies and showed that genetic stimulation of autophagy increases proliferation and restrains other hallmarks of aging, such as mitochondrial dysfunction, loss of proteostasis, and cellular senescence [11,38,47-49]. In light of evidence that autophagy declines with age in osteoblast lineage cells [6-8], and that loss of autophagy mimics several aspects of skeletal aging [1], we propose that declining autophagy could be contributing to age-related bone loss. If this is the case, stimulation of autophagy may blunt age-related bone loss. To test this, others have administered rapamycin to old mice (16- and 20-month-old) [6,50] and rats (24-month-old) [51] with varying doses and intervals. While two of these studies show a beneficial impact on trabecular bone, the other did not observe any effect on trabecular or cortical bone. Part of the discrepancy between these studies may be due to differences in dose and interval of administration, or age or sex of the model organism. However, as stated above, all three studies are confounded by the lack of cellular and molecular specificity of this mTOR inhibitor. Thus, *Tfeb*^CRa^ mice will also be a valuable tool for determining if declining autophagy in cells of the osteoblast lineage is sufficient to prevent any aspects of skeletal aging.

In conclusion, increasing endogenous *Tfeb* expression in the osteoblast lineage promotes autophagy, and bone formation, leading to a remarkable increase in bone mass. These findings lay the groundwork for future studies aimed at examining whether stimulation of autophagy can blunt skeletal pathologies associated with insufficient autophagy or decreased bone formation.

## METHODS

### Testing *Tfeb* sgRNAs in vitro

Five candidate sgRNAs were identified using the Zhang Laboratory CRISPR Design tool (http://crispr.mit.edu) by searching the region 150 bp upstream and downstream from the *Tfeb* TSS. sgRNAs with high-quality scores, minimal off-target scores, and with no potential off-targets in other genes were chosen as candidates. pX330 [52] (Addgene plasmid #42230) and pX458 [53] (Addgene plasmid #48138) were obtained from the Zhang Laboratory via Addgene (Cambridge, MA, USA). The Cas9 sequence of the pX330 plasmid was replaced with eGFP cloned from the px458 plasmid producing the sgRNA_GFP plasmid. Each of the five candidate *Tfeb* target sgRNA sequences indicated below was cloned into a sgRNA_GFP plasmid producing *Tfeb* sgRNA 1 through 5. Sequences for the sgRNAs are as follows: sgRNA-1: AAATCCCGGCGAGCCCTTCG (PAM: CGG), sgRNA-2: ACATTTCCCAGCGGGCACAG (PAM: CGG), sgRNA-3: TGCCGGGCGCAGCGGGAAGT (PAM: GGG), sgRNA-4: AACGGAGACGGCGCCGACAG (PAM: CGG), sgRNA-5: CGGGACGCAGAGAACGGAGA (PAM: CGG). Each *Tfeb* sgRNA plasmid was co-transfected with SP-dCas9-VPR plasmid [18] (a gift from George Church, Addgene plasmid # 63798) into UAMS-32 cells using PolyJet reagent (SignaGen Laboratories, Gaithersburg, MD, USA) according to the manufacturer’s instructions. After 48 hours, transfected cells were subjected to fluorescence-activated cell sorting (FACS) to acquire GFP-positive cells, which contain CRISPRa components, and sorted into 3 wells of a 96-well plate at a density of 7,500 cells per well. Cells were incubated overnight and then, Cells-to-Ct kit (Invitrogen, A25605) was used to harvest RNA and measure gene expression via qRT-PCR.

### Generation of Mice

The sgRNA*^Tfeb^*mice were produced via CRISPR/Cas9-mediated knock-in of a U6-sgRNA*^Tfeb^*_4 cassette into the Rosa26 locus as previously described [22,54]. Briefly, sgRNA*^Tfeb^*knock-in mice were produced by microinjection of 50 ng/μl Cas9 protein, 30 ng/μl sgRNA*^Rosa^*, and 10 ng/μl single-strand oligonucleotide donor (ssOND) into the pronuclei of C57BL/6J mice. sgRNA*^Rosa^* (ACTCCAGTCTTTCTAGAAGA, PAM: TGG) is expected to facilitate double-strand break production by the Cas9 protein at the 1^st^ intron of the Rosa loci [54]. The ssOND containing the sequence of U6-*Tfeb*_sgRNA_4 is expected to work as a template for homologous recombination-mediated repair of this double-strand break, thereby leading to the insertion of the U6-*Tfeb*_sgRNA_4 cassette into the Rosa locus. Founders were screened for the presence of the knock-in sequence using the following primers: F1 5’-AAGCACTTGCTCTCCCAAAG-3’; R1 5’-GGCGGATCACAAGCAATAAT-3’. This PCR produced an 803bp band for the knock-in allele and a 447bp band for the wild-type allele. The knock-in was confirmed by DNA sequencing of the PCR product. The progeny of the founders were genotyped using genomic DNA isolated from the tail tips with primers: F1 and R1 primers indicated above, and F2 5’-GAGGGCCTATTTCCCATGAT-3’, and R2 5’-GGTGTTTCGTCCTTTCCACA-3’ primers which produce a 210 bp PCR band for the knock-in allele. CRa [19] and Osx1-Cre [23] mice were previously produced by Zhou et al. and Rodda et al., respectively. These mice were obtained from and genotyped according to the directions of the Jackson Laboratory (Strain #: 0331645 and Strain #:006361). All mice were provided food and water *ad libitum* and were maintained on a 12-hour light/dark cycle. All animal studies were carried out with approval from and following the policies of the Institutional Animal Care and Use Committee of the University of Arkansas for Medical Sciences. The studies described here were performed and reported in accordance with ARRIVE guidelines. Each figure legend indicates the sex, number, and age of the mice used for each experiment.

### Ex vivo osteoblastic cell culture

Bone marrow stromal cells were flushed from the tibias and femurs of mice using an isolation medium (Hanks Buffered Saline containing 10% fetal bovine serum and 1% Penicillin/Streptomycin/Glutamine). The cells were centrifuged, transferred into osteogenic medium (α-MEM containing 10% fetal bovine serum, 1% penicillin/streptomycin/glutamine, and 50 μg/ml ascorbic acid), filtered through a 75 µm cell strainer to obtain a single cell suspension, and then seeded into a 12-well-plate at 5×10^6^ cells/well. Half of the culture media was changed every 2 to 3 days for 3 weeks. After 21 days, protein were isolated for downstream analysis.

For ex vivo mineral deposition assay, cells were isolated as indicated above and then plated at a density of 2 ×10^6^ cells/well in osteogenic media with 10mM β-glycerolphosphate. Half of the culture media was changed every 2 to 3 days for 3 weeks. The cultures were then fixed with 50% ethanol at 4 °C for 15 minutes, dried for 2 hours, and stained with an aqueous solution of 40 mM alizarin red for 30 min at room temperature.

### Ex vivo proliferation assay

Bone marrow stromal cells were flushed from the tibias and femurs of mice and red blood cells were removed using ACK buffer (0.01 mM EDTA, 0.011 M KHCO3, and 0.155 M NH4Cl, pH 7.3). The remaining cells were cultured with osteogenic media (10% FBS, 1% PS, and 50 μg/ml ascorbic acid). After cells reached 80% confluence (5 to 10 days), cells were re-plated.

To measure cell proliferation, collagen working solution (Sigma, C3867) was prepared in PBS (10 μg/mL). Working solution was added to each well of the 8-well Lab-Tek™ Glass Chamber Slide System, incubated for 2 hours at 37 °C and 5 % CO_2_, and washed once with PBS. Cells were plated in 96-well cell-culture plates at 5×10^4^ cells/cm^2^, in medium with 10% or 1% FBS. After 24 hours, medium was replaced with medium containing 100 μM BrdU. The plates were then incubated at 37 °C and 5 % CO2 for 24 or 48 hours. The cell proliferation assay was performed according to the manufacturer’s instructions (Roche Applied Sciences, 11647229001).

### Lysotracker staining of osteoblasts ex vivo

Bone marrow stromal cells were flushed from the tibias and femurs of mice and red blood cells were removed using ACK buffer (0.01 mM EDTA, 0.011 M KHCO3, and 0.155 M NH4Cl, pH 7.3). The remaining cells were cultured with osteogenic media (10% FBS, 1% PS, and 50 μg/ml ascorbic acid). After cells reached 80% confluence (5 to 10 days), cells were re-plated.

For flow cytometry, cells were plated in 6-well plates at 2.5×10^4^ cells/cm^2^. After 24 hours, cells were incubated with 50 nM of Lysotracker (Invitrogen, L12492) for 30 min, in fresh cell culture medium with no FBS at 37 °C and 5 % CO_2_. After treatment, cells were washed with PBS, resuspended in Ca/Mg^2+^ free PBS with 0.5% BSA, and analyzed by flow cytometry. All samples were acquired using BD LSRFortessa (BD Biosciences, San Jose, California, USA) with FACSDiva™ software version 6.1.3. Data was analyzed using FlowJo 10.10 (Tree Star Inc, Ashland, USA). The cell populations were first plotted according to their size and complexity on a Forward Scatter-A (FSC-A) vs. Side Scatter-A (SSC-A) plot, to remove any dying cells and debris, followed by FSC-A vs. FSC-H plot to isolate single cells. The median fluorescent intensity (MFI) of Lysotracker was determined using the APC filter.

### RNA isolation and gene expression analysis

Lumbar vertebrae 5, tibia shafts, and calvaria were dissected and cleaned of soft tissue. The tibia epiphyses were removed, and the marrow was flushed from the shafts. The bones were snap-frozen in liquid nitrogen and stored at −80°C. For RNA isolation, the bones were homogenized in Trizol Reagent (Life Technologies, 15596018). Next, RNA was isolated from homogenized bones with the RNAeasy Plus Mini Kit (Qiagen, 74136) according to the manufacturer’s instructions. RNA concentrations were determined with a Nanodrop instrument (Thermo Fisher Scientific). 1 μg RNA was used to synthesize cDNA with a High-Capacity cDNA Reverse Transcription Kit (Applied Biosystems, 4368814). Relative abundance of mRNAs were measured using multiplex quantitative real-time PCR (qRT-PCR) with TaqMan Fast Advanced Master Mix (Applied Biosystems, 4444964), FAM-labeled TaqMan gene expression assays (Life Technologies), and VIC-labeled mouse Actb (β-actin) (Applied Biosystems, 4352341E). The following FAM-labeled assays were used in the gene expression analysis: *Lamp1* (Mm01217070_m1), *CtsB* (Mm01310506_m1), *CtsD* (Mm00515586_m1), *CtsF* (Mm00490782_m1), *Clcn7* (Mm00442400_m1), *Gns* (Mm00659592_m1), *Hexa* (Mm00599877_m1), *Mcoln1* (Mm00522550_m1), *Tpp1* (Mm00487016_m1), *Map1lc3* (LC3, Mm00782868_sH), *Atg7* (Mm00512209_m1), *Becn1* (Mm01265461_m1), *Parkin* (Mm01323528_m1), *Pink* (Mm00550827_m1), *Bnip3* (Mm01275600_g1), *Bnip3l* (Mm00786306_s1), *Mfn2* (Mm00500120_m1), *Fundc1* (Mm00511132_m1), *Sp7* (Osx, Mm04209856_m1), *Col1a1* (Mm00801666_g1), *Acp5* (Mm00475698_m1), *CtsK* (Mm00484039_m1), *Bglap* (OCN, custom). The relative mRNA levels were calculated using the comparative cycle threshold (ΔCt) method [55].

### Immunoblot analysis

Protein was extracted from cultured cells using RIPA Buffer (Fisher Scientific, PI89901) with protease/phosphatase inhibitors (Cell Signaling Technologies, 5872S) according to the manufacturer’s instructions. Protein was extracted from humeri shafts previously snap-frozen in liquid nitrogen and stored at −80°C. Once pulverized, bone pieces were collected into RIPA Buffer with protease/phosphatase inhibitors. Proteins were resolved in 4–20% or 4–15% Mini-PROTEAN TGX gels (BIORAD, 4561093 and 4561083, respectively) and transferred onto TransblotTurbo midi-size nitrocellulose membranes (0.2 μm pore size, BIORAD, 1704271). The membranes were blocked for 30 min with LI-COR Blocking Buffer-PBS (LI-COR, 4561083) and incubated overnight with primary antibodies, rocking at 4°C. The primary antibodies used were TFEB (Invitrogen, PA5-96632, 1:500 dilution), LAMP1 (Cell Signaling Technology, 99437, 1:1000 dilution), p62 (Cell Signaling Technology, 23214s, 1:1000 dilution), LC3 (Cell Signaling Technology, 12741T, 1:1000 dilution), Sclerostin (R&D Systems, AF1589, 1:500 dilution) and β-actin (Millipore Sigma, A5316, 1:4000 dilution). After overnight incubation, membranes were washed three times with PBS for 5 minutes each and incubated for 45 min with appropriate secondary antibodies conjugated with IRDye 680 or IRDye 800 dyes (LI-COR, 1:2000 dilution). After washing with PBS three times 5 minutes each time, membranes were scanned and analyzed with an Odyssey IR imaging system (LI-COR) and Image Studio Software (Version 5.2).

### Skeletal phenotyping

Bone mineral density (BMD) of the total body, spine, and femur was measured by dual-energy x-ray absorptiometry (DXA) with a PIXImus Mouse Densitometer (GE Lunar Corp., Madison, WI) as previously described [56]. Calvaria and shoulder blades were excluded from the analysis of total body BMD. Vertebral BMD was measured in lumbar vertebrae 2-6, and femoral BMD was measured using the right femur.

For µCT analysis, lumbar vertebrae 5, femurs, and humeri were dissected, cleaned of soft tissue, wrapped in saline-soaked gauze, and stored at −20°C. Samples were thawed before scanning and medium-resolution scans were obtained (12 μm isotropic voxel size) using a model uCT40 (Scanco Biomedical) as previously described [22]. A Gaussian filter (sigma = 0.8, support = 1) was used to reduce noise and a threshold of 220 was used for all scans. Long bone midshaft cortical measurements were performed by drawing contours to measure the cortical thickness on the 20 slices flanking the midshaft. Long bone trabecular analysis was performed by drawing contours every 20 slices of the distal metaphysis beginning 10 slices proximal to the growth plate and ending 151 slices proximal to the first contour. Vertebral cortical thickness and trabecular analyses were performed by drawing contours every 10 slices between the two growth plates of the vertebrae. The vertebral cortical bone thickness was determined on the ventral cortical wall using contours of cross-sectional images, drawn to exclude trabecular bone.

### Biomechanical testing

The mechanical properties of the femur were determined by a three-point bending analysis using ElectroForce 5500 (TA Instruments, New Castle, DE) as previously described [57]. Femurs were placed posterior-side down on a miniature bending apparatus with lower supports set at 8 mm apart with the left support set proximal to the distal condyles. The test was conducted at room temperature. Load was applied to the anterior surface of the femur midway between the lower supports. The load was applied at a constant rate of 3 mm/min until failure. Structural or extrinsic properties (stiffness, ultimate force, and energy to ultimate force) and material or intrinsic properties (ultimate stress, toughness to ultimate load, and Young’s Modulus) were derived by measuring the maximum load and displacement of the femur and normalized for bone size. The external measurements (length, width, and thickness) of the femora were recorded with a digital caliper. Bone geometry and volume were determined by µCT.

### Histology

Mice were injected with 20 mg/kg of calcein (Sigma Aldrich, C0875) and alizarin complexone (Sigma Aldrich, A3882), 6 days and 2 days prior to euthanization, respectively. For histology, femurs or L1-3 vertebrae were fixed in 10% Millonig’s formalin for 24 hours, and then dehydrated using a series of serial dilutions of ethanol into 100% ethanol. Next, the bones were embedded in methyl methacrylate, and longitudinal sections were obtained. To measure bone formation at the periosteal, endosteal, and trabecular surfaces, dynamic histomorphometry was performed on the unstained sections. Mineralizing surface and mineral apposition rate were quantified using the Osteomeasure system (OsteoMetrics, Decatur, GA) interfaced to an Axio image M2 (Carl Zeiss, NY), and bone formation rate was calculated from these values. Results were reported using terminology recommended by the Histomorphometry Nomenclature Committee of the American Society for Bone and Mineral Research [58].

### Serum bone turnover assays

Blood was collected from the facial vein and incubated at room temperature for a minimum of 30 minutes to allow the blood to clot. Serum was collected by serial centrifugation (twice 2,000 g for 12 minutes) and stored at −20°C. Procollagen type 1 N-terminal propeptide (P1NP) and tartrate-resistant acid phosphatase form 5b (TRAcP5b) were measured following the manufacturer’s instructions (Immunodiagnostic Systems, AC-33F1 and SB-TR103, respectively). Circulating sclerostin levels were measured using Mouse/Rat SOST/Sclerostin Quantikine ELISA Kit (R&D Systems, MSST00).

### Statistical analysis

All values are reported as mean ± standard deviation (SD). Differences between the Cre-expressing controls (Osx1-Cre, Osx1-Cre;CRa, Osx1-Cre;sgRNA*^Tfeb^*) and *Tfeb*^CRa^ (Osx1-Cre;CRa;sgRNA*^Tfeb^*) mice were evaluated using GraphPad Prism 7.05 software (GraphPad Software, Inc, La Jolla, CA, USA). The specific statistical tests and number of replicates performed are indicated in figure legends. For Figure 6, a repeated measures model was fit to BMD measurements focusing on the effects of group and month, along with the interaction of group and month and with an autoregressive (1) covariance structure using SAS version 9.4. Residuals of the model were tested for normality and constant variance.

## ACKNOWLEDGEMENTS

We thank the Genetic Models Core Facility for the production of the sgRNA*^Tfeb^* murine model, the Bone Histology and Imaging Core for their help with tissue collection and analysis, and the staff of the UAMS Department of Laboratory Animal Medicine for their help with husbandry and care of mice. A.Y.S. received salary support by T32-AR065971 and the National Center for Advancing Translational Sciences of the National Institutes of Health under Award Numbers KL2TR003108 and UL1TR003107.

## DATA AVAILABILITY STATEMENT

The data that support the findings of this study are contained in the manuscript and its Supplementary Materials files. The raw data are available from the corresponding authors upon reasonable request.

---

**Figure S1.**
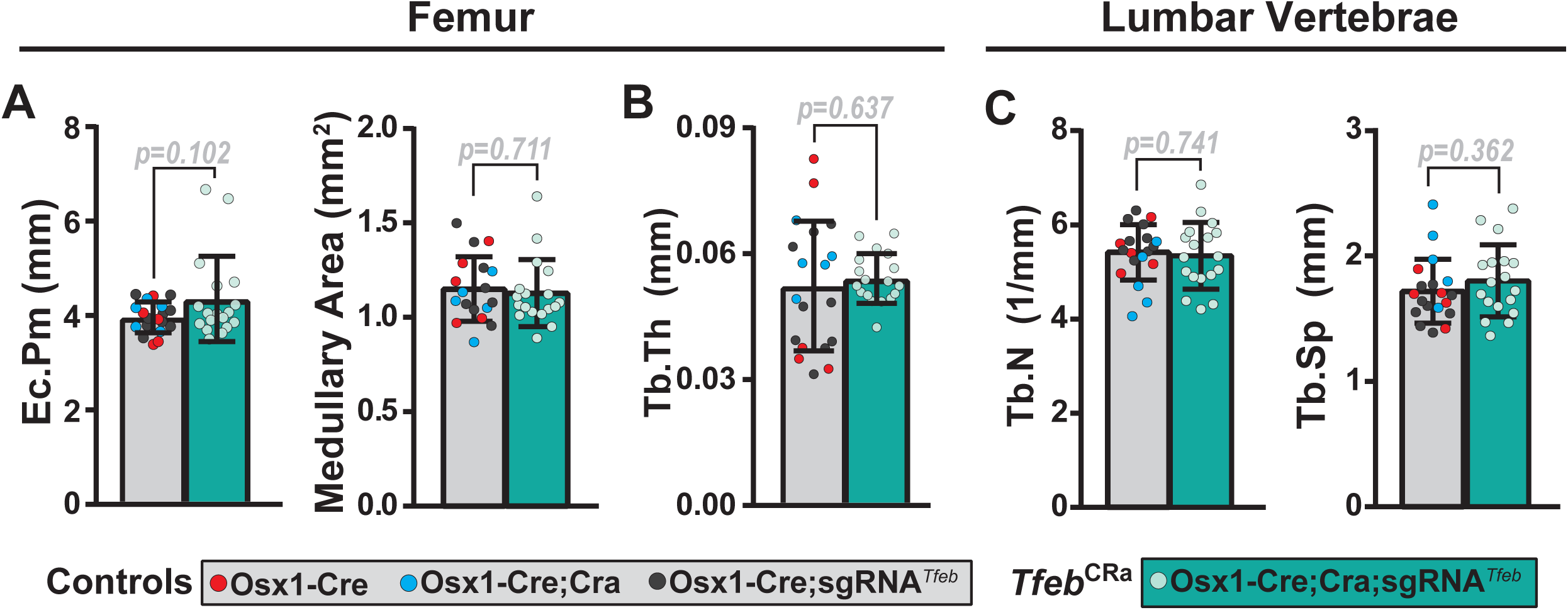
*Tfeb* elevation does not alter femur endosteal perimeter or vertebral trabecular number and spacing at 4.5 months of age. (**A-C**) µCT analysis was performed on femurs and lumbar vertebrae 4 (L4) from 4.5-month-old male *Tfeb*^CRa^ mice and their littermate controls. (**A**) Endosteal perimeter (Ec.Pm) and medullary area measured at the femur midshaft. (**B**) Trabecular thickness (Tb.Th) measured at the femoral distal metaphysis. (**C**) Trabecular number (Tb.N) and trabecular spacing (Tb.Sp) measured in L4. Bars indicate mean ± SD. n = 17-19 mice/group. Indicated *p* values were calculated by unpaired *t*-test.

**Figure S2.**
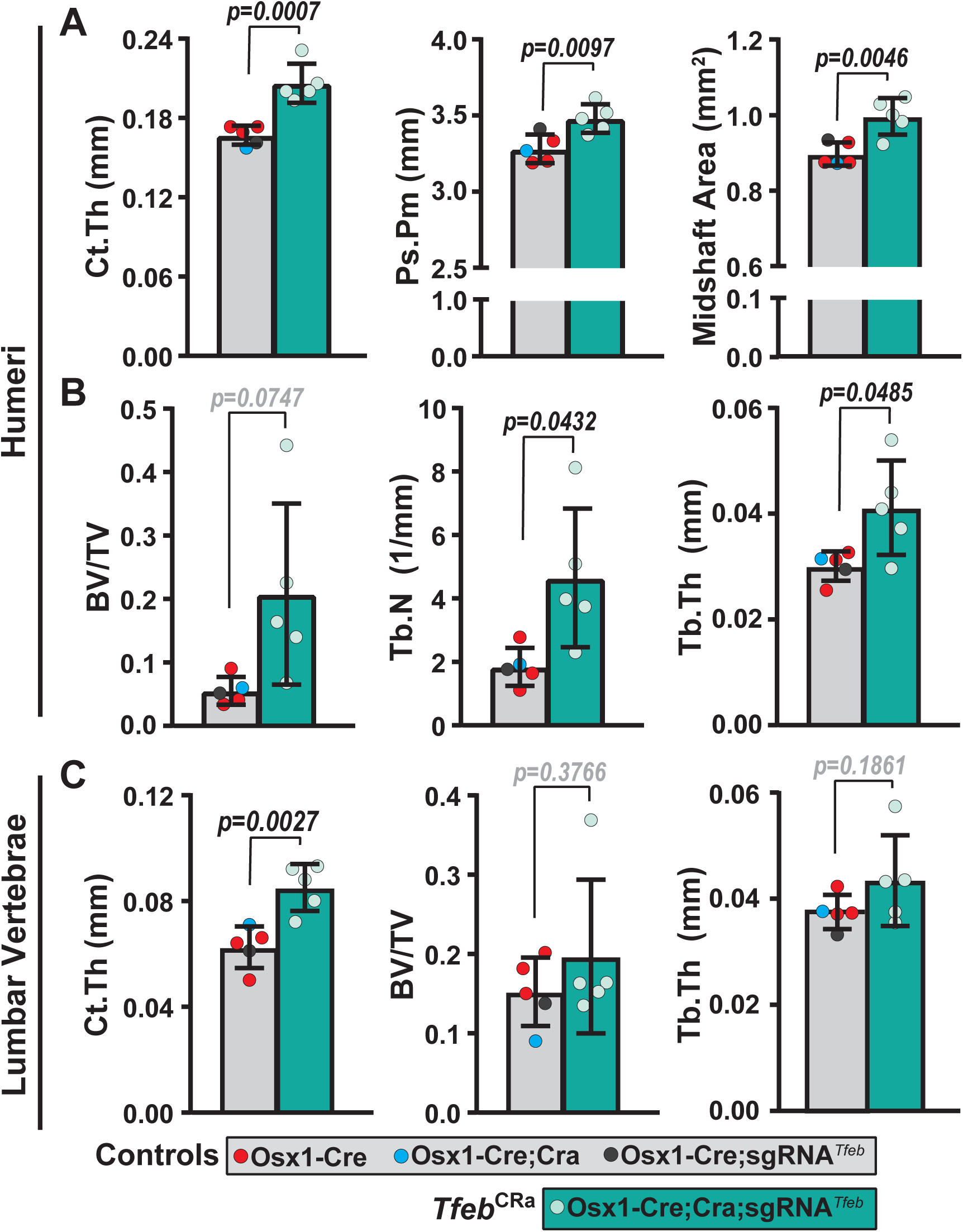
*Tfeb* elevation increases bone mass in 4.5-month-old female mice. µCT analysis was performed on humeri and lumbar vertebrae 4 (L4) from 4.5-month-old female *Tfeb*^CRa^ mice and their littermate controls. (**A**) Cortical thickness (Ct.Th), periosteal perimeter (Ps.Pm), and midshaft area measured at the humoral midshaft. (**B**) Trabecular bone volume over tissue volume (BV/TV), trabecular number (Tb.N), and trabecular thickness (Tb.Th) were measured at the humoral distal metaphysis. (**C**) Ct.Th, BV/TV, and Tb.Th measured in L4. Bars indicate mean ± SD. n = 5 mice/group. Indicated *p* values were calculated by unpaired *t*-test.

**Figure S3.**
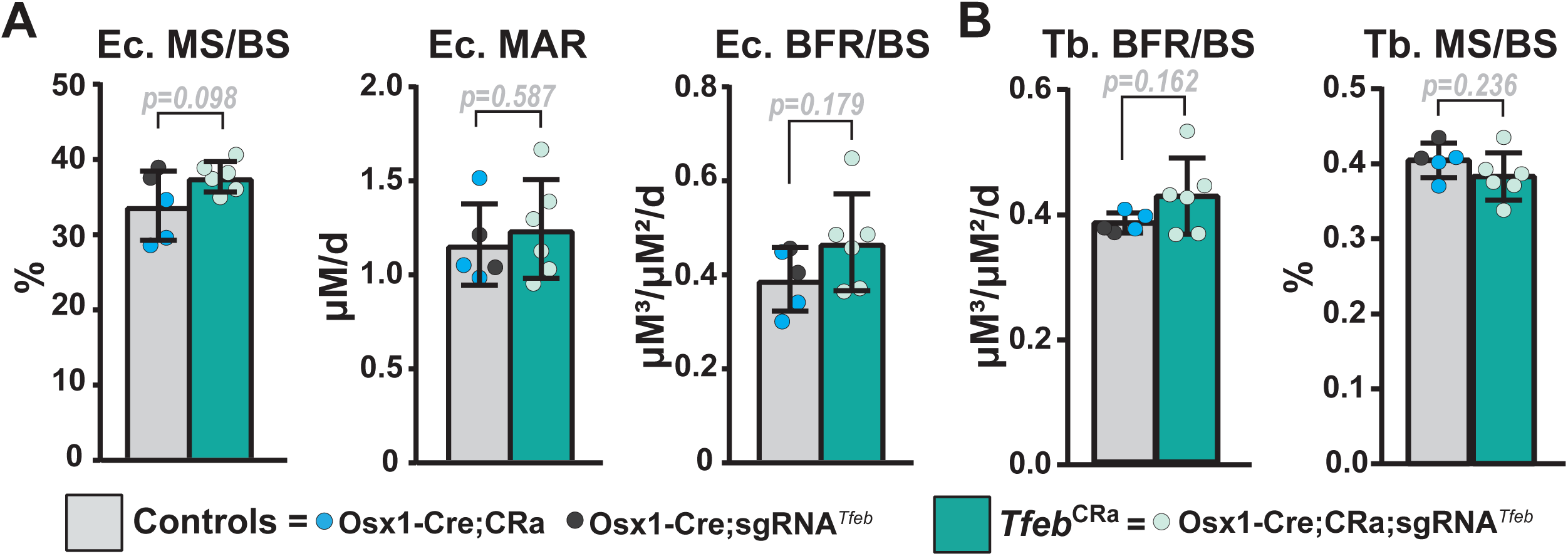
*Tfeb* elevation does not change bone formation at the femoral endosteal and trabecular surfaces at 4.5 months of age. Dynamic histomorphometric analysis was performed on femurs of 4.5-month-old male *Tfeb*^CRa^ mice and their littermate Cre controls. (**A**) Quantification of the mineralizing surface per bone surface (MS/BS), mineral apposition rate (MAR), and bone formation rate per bone surface (BFR/BS) at the endosteal surface. (**B**) Quantification of BFR/BS and MS/BS at the trabecular surface. Bars indicate mean ± SD. n = 5-6 mice/group. Indicated *p* values were calculated by unpaired *t*-test.

**Figure S4.**
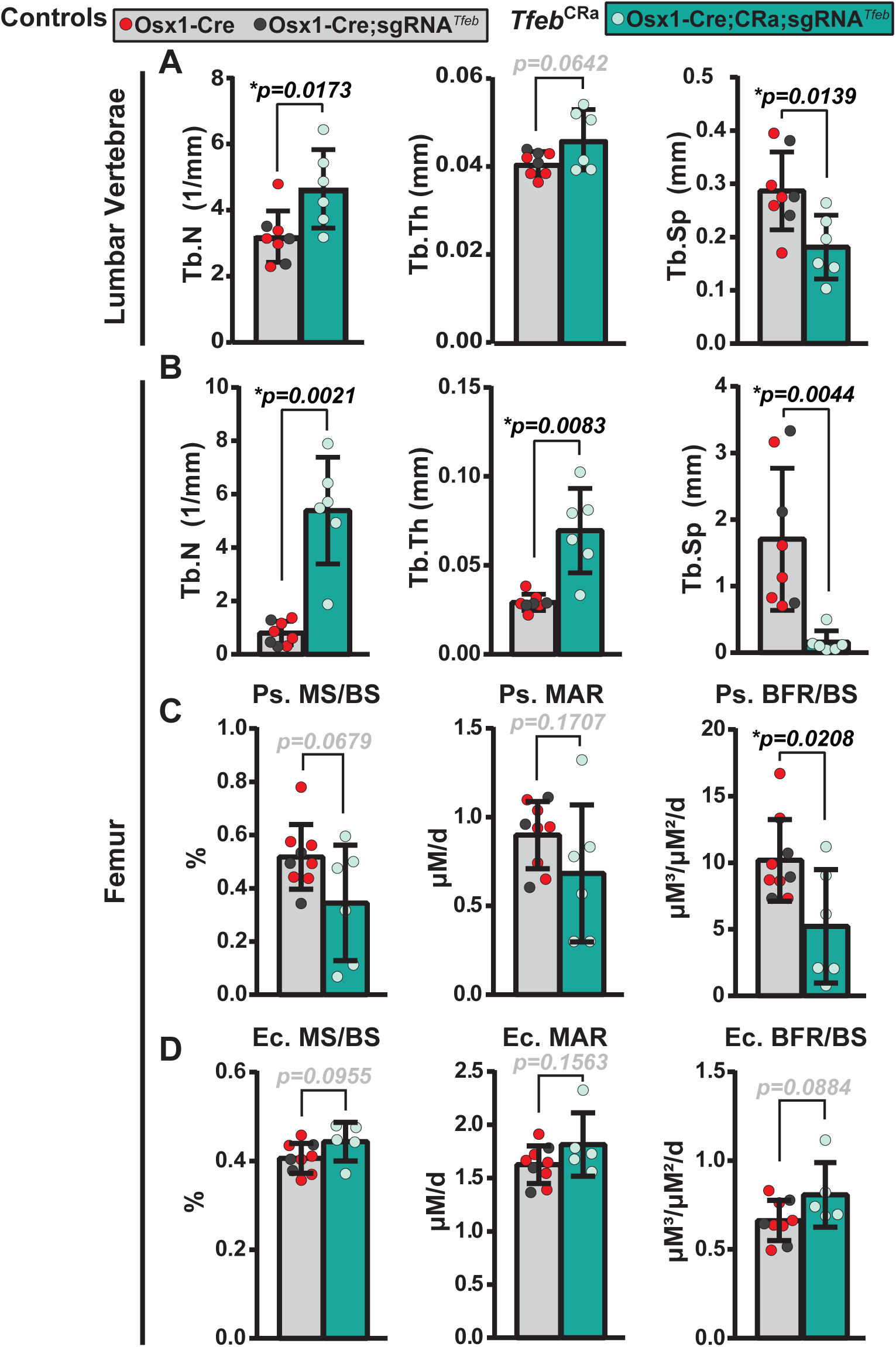
*Tfeb* elevation is anabolic up to 12 months of age. (**A** and **B**) µCT analysis was performed on lumbar vertebrae 4 (L4) and femurs from 12-month-old female *Tfeb*^CRa^ mice and their littermate controls. (**A** and **B**) Trabecular number (Tb.N), trabecular thickness (Tb.Th), and trabecular spacing (Tb.Sp) measured in L4 (**A**) and at the femoral distal metaphysis (**B**). (**C** and **D**) Dynamic histomorphometry was performed on femurs of 12-month-old female *Tfeb*^CRa^ mice and their littermate Cre controls. (**C** and **D**) Quantification of the mineralizing surface per bone surface (MS/BS), mineral apposition rate (MAR), and bone formation rate per bone surface (BFR/BS) at the femoral periosteal (**C**) and endosteal (**D**) surfaces. Bars indicate mean ± SD. n = 5-9 mice/group. Indicated *p* values were calculated by unpaired *t*-test.

**Figure S5.**
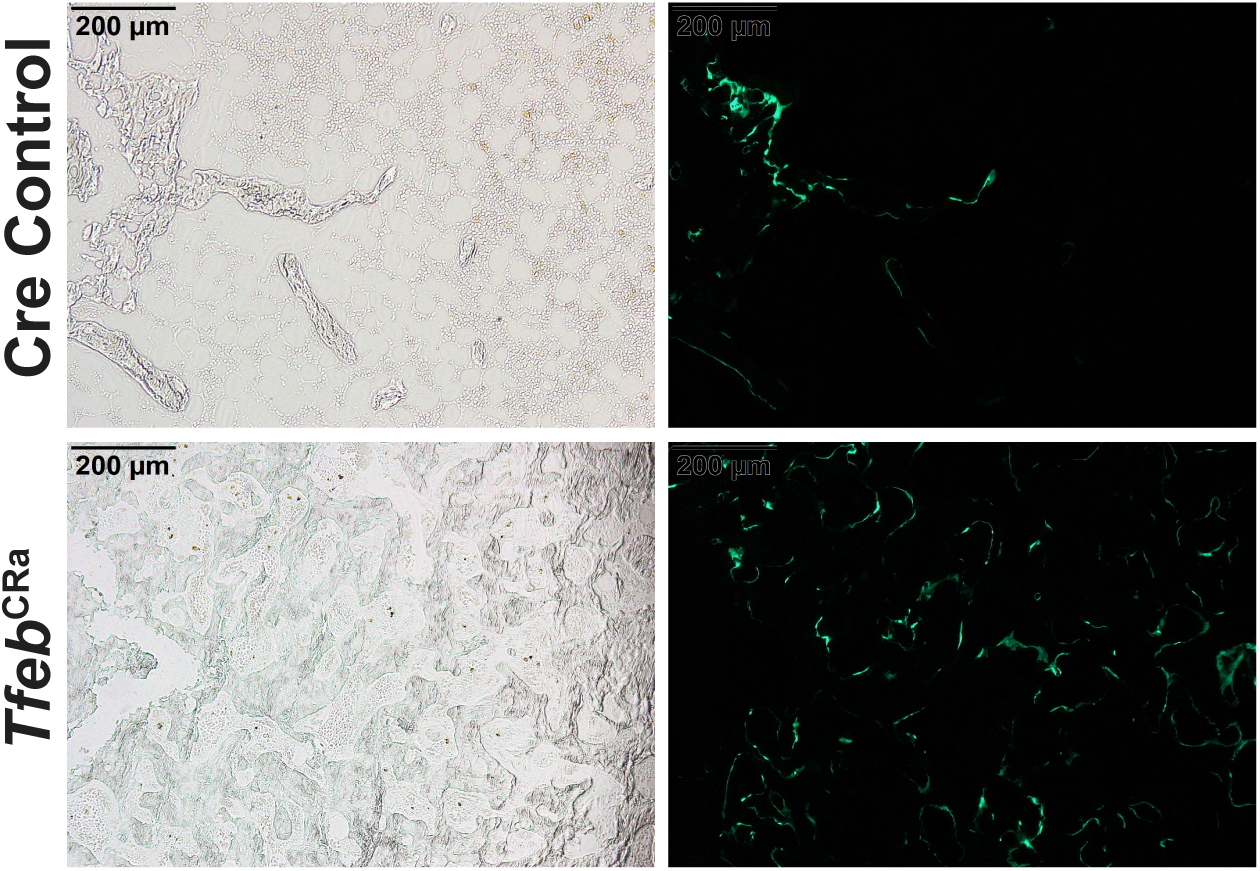
*Tfeb* OE increases bone formation at the femoral trabecular bone at 12 months of age. Light microscopy (left panel) and fluorescent microscopy (GFP filter, right panel) images of femoral trabecular bone of 12-month-old female *Tfeb*^CRa^ and control mice.

**Figure S6.**
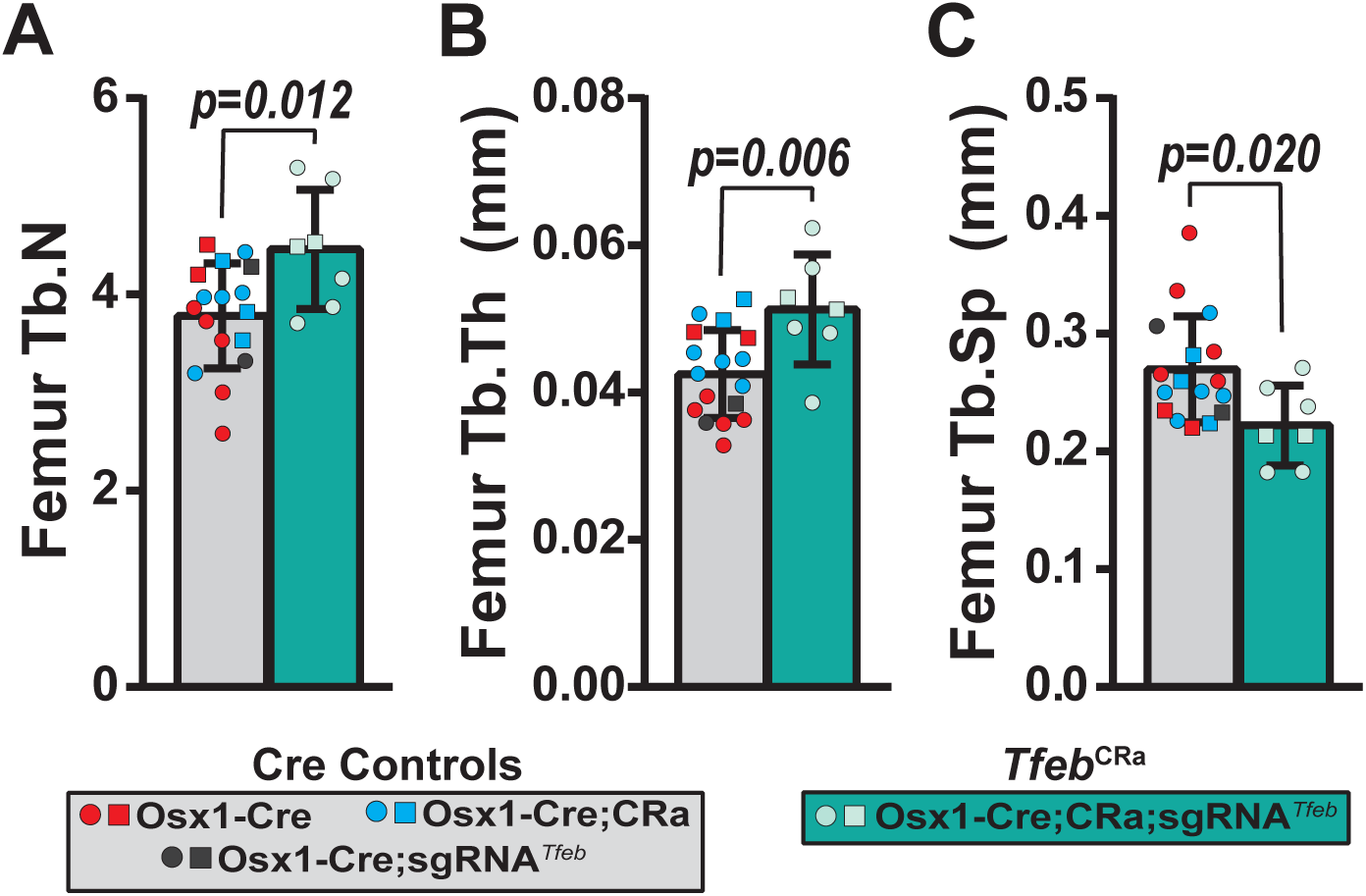
*Tfeb* elevation in the osteoblast lineage increases femoral trabecular bone at 8 weeks of age. (**A-C**) µCT analysis of the femoral distal metaphysis of 2-month-old male (square) and female (circle) *Tfeb*^CRa^ mice and their littermate controls to quantify trabecular number (Tb.N) (**A**), trabecular thickness (Tb.Th) (**B**) and trabecular spacing (Tb.Sp) (**C**). Bars indicate mean ± SD. n = 7-17 mice/group. Indicated *p* values were calculated by unpaired *t*-test. Unpaired *t*-test with Welch’s correction was used if the SD of groups were different.

